## Supplementary material for "*DNAJB1-PRKACA* in HEK293T cells induces *LINC00473* overexpression that depends on PKA signaling"

**Corresponding Author:**

Khashayar Vakili, MD

Supplementary Figure S1.

(a)

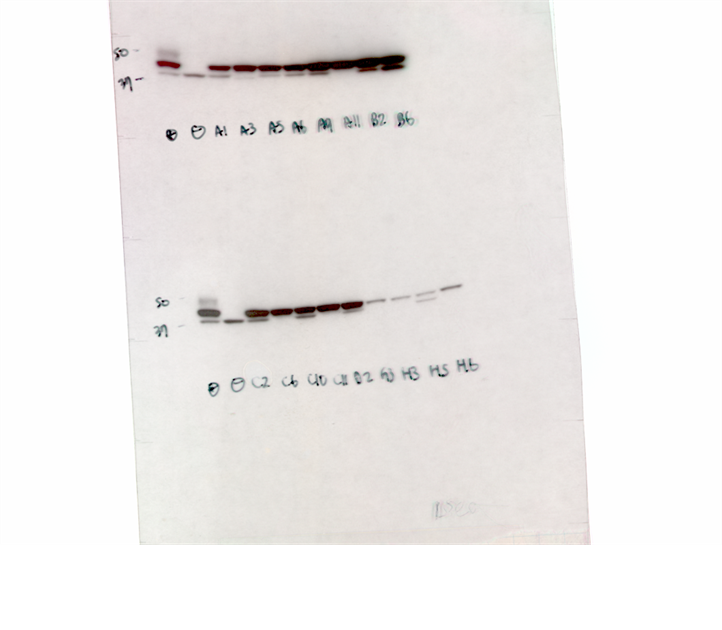

(b)

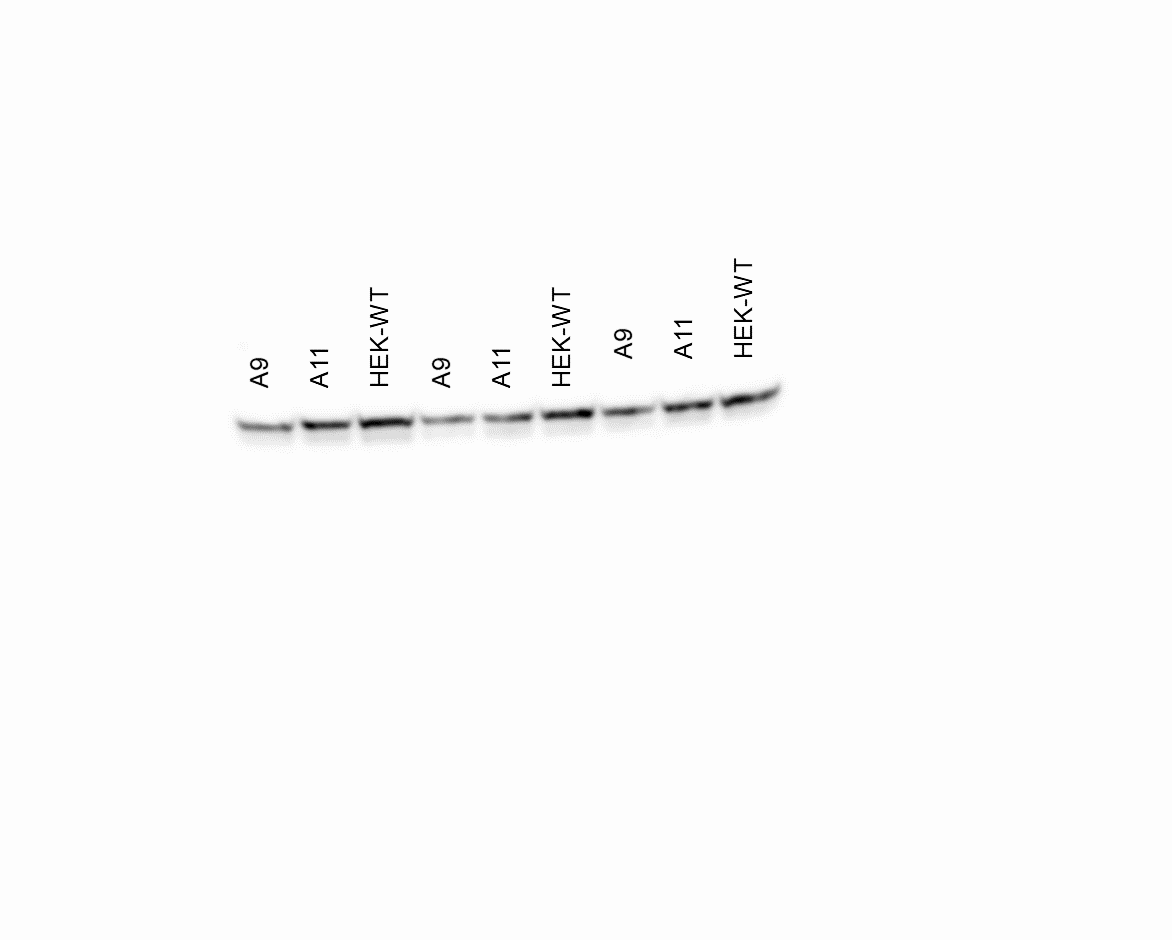

(c)

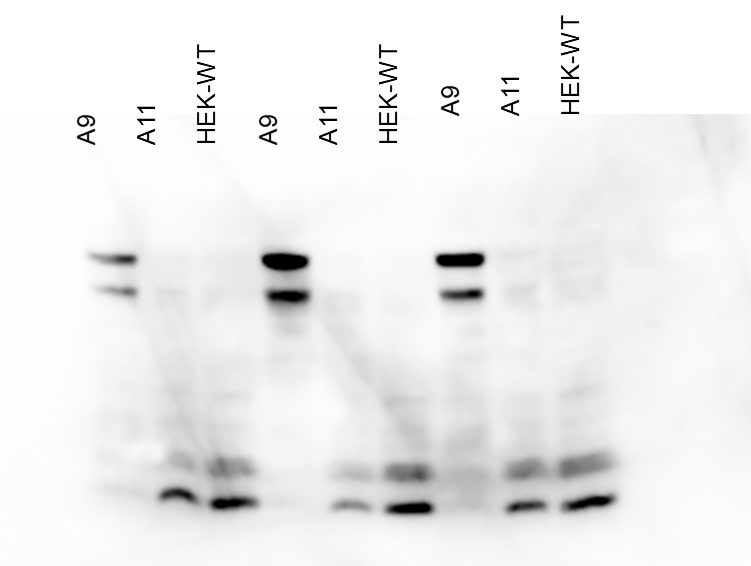

(d)

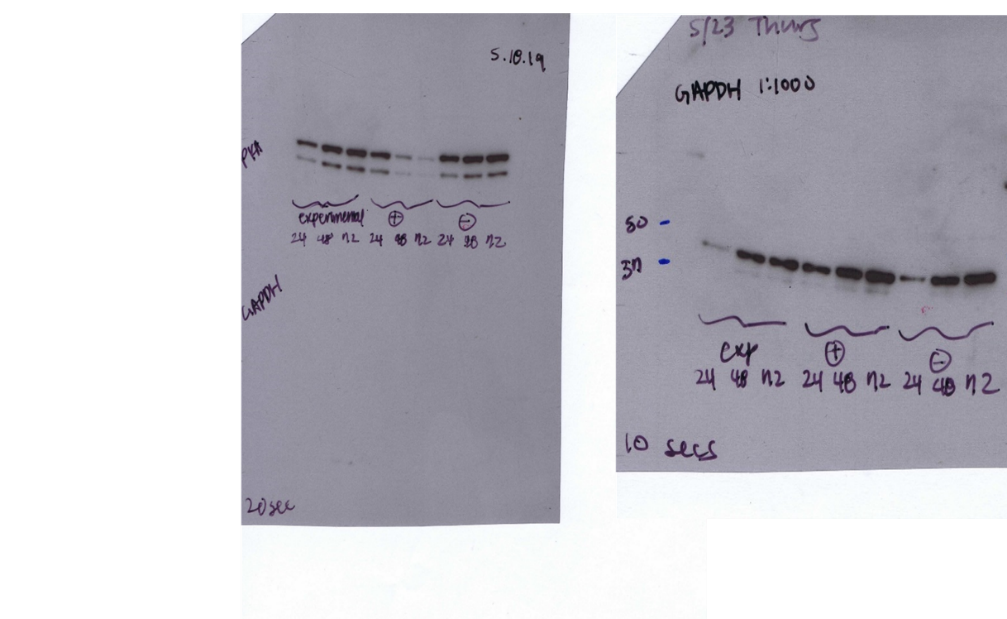

**Supplementary Figure S1.** **Full-length immunoblots.** (a) Blots demonstrate the expression of PKA-Cα and DP fusion protein in engineered HEK-DP clones as presented in Figure 1. (b) CREB immunoblot in A9, A11, and HEK-WT performed in triplicate. (c) Phosphorylated CREB in A9, A11, and HEK-WT. (d) α−PKA immunoblot of A9 cells following siRNA treatment (right panel) and corresponding GAPDH immunoblot (left panel).

Supplementary Figure S2.

(a)

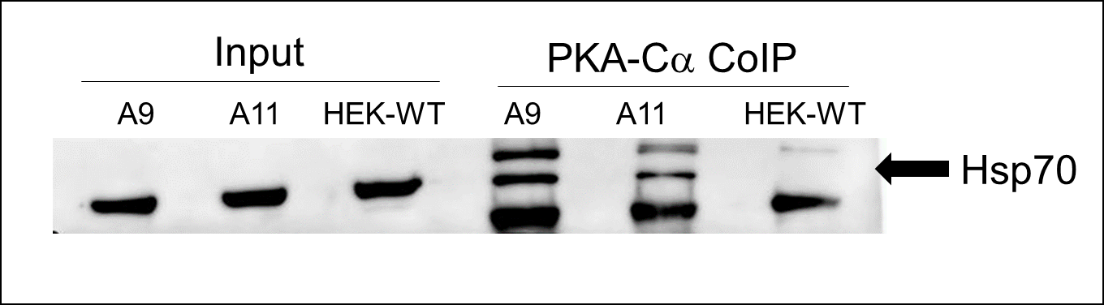

(b)

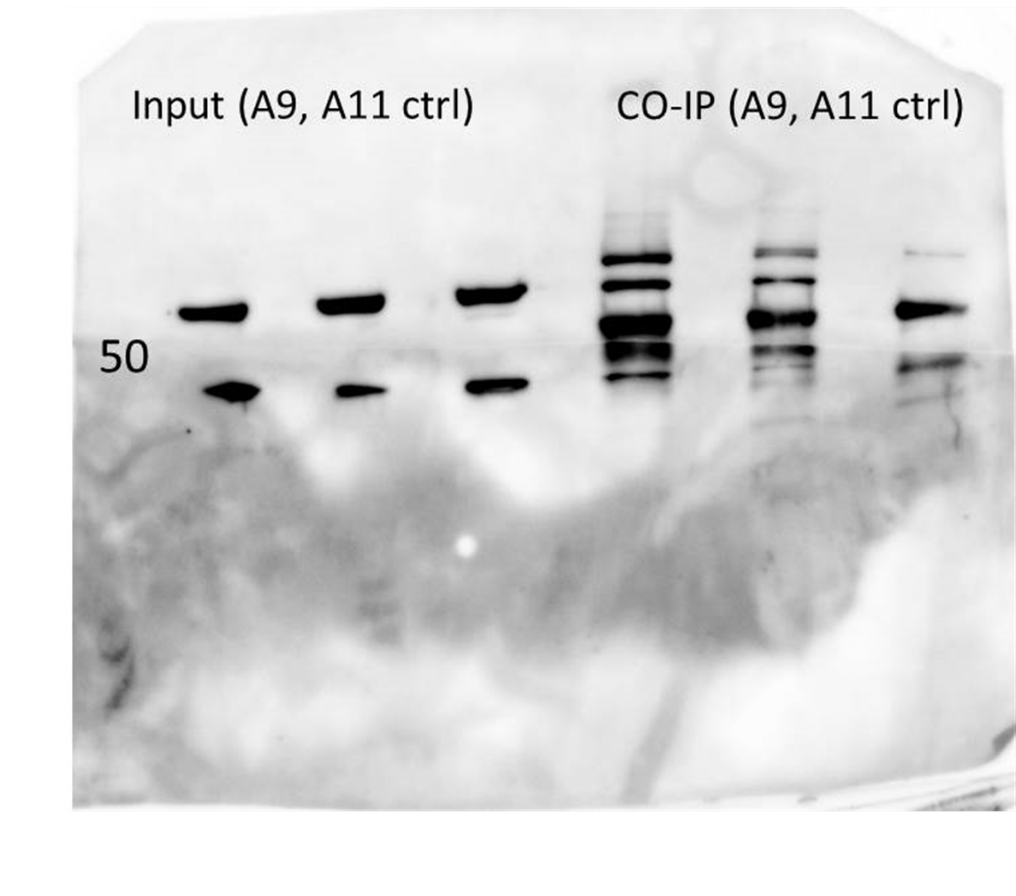

**Supplementary Figure S2.** **Hsp70 interacts with PKA-Cα in A9 and A11 cells.** (a) Hsp70 expression is demonstrated in all cell lines. Following co-immunoprecipitation with PKA-Cα antibody, Hsp70 is identified in the protein complexes from A9 and A11. (b) Full-length blot represented in panel A.

Supplementary Table S1.

| **Gene** | **Protein** | **A9 vs. HEK-WT**  **Adj. p-value** | **A11 vs. HEK-WT**  **Adj. p-value** |
| --- | --- | --- | --- |
| DNAJB1 | DnaJ homolog subfamily B member 1 | 3.09E-06 | 8.73E-07 |
| CTSH | Pro-cathepsin H | 3.09E-06 | 8.73E-07 |
| PRKACA | cAMP-dependent protein kinase catalytic subunit alpha | 3.09E-06 | 8.73E-07 |
| RSRC2 | Arginine/serine-rich coiled-coil protein 2 | 3.09E-06 | 8.73E-07 |
| XRN2 | 5-3 exoribonuclease 2 | 3.09E-06 | 8.73E-07 |
| RPL9 | 60S ribosomal protein L9 | 3.09E-06 | 8.73E-07 |
| GANAB | Neutral alpha-glucosidase AB | 3.09E-06 | 8.87E-07 |
| ACSL3 | Long-chain-fatty-acid--CoA ligase 3 | 3.26E-06 | 8.73E-07 |
| EMC7 | ER membrane protein complex subunit 7 | 3.32E-06 | 8.73E-07 |
| MYO1C | Unconventional myosin-Ic | 3.32E-06 | 8.73E-07 |
| CFL1 | Cofilin-1;Cofilin-2 | 3.32E-06 | 8.73E-07 |
| LIMA1 | LIM domain and actin-binding protein 1 | 3.32E-06 | 8.73E-07 |
| PFDN2 | Prefoldin subunit 2 | 3.32E-06 | 8.73E-07 |
| AAAS | Aladin | 3.32E-06 | 9.47E-07 |
| BAG2 | BAG family molecular chaperone regulator 2 | 3.4E-06 | 1.12E-06 |
| LUC7L3 | Luc7-like protein 3 | 3.5E-06 | 1.04E-06 |
| LRPPRC | Leucine-rich PPR motif-containing protein, mitochondrial | 3.65E-06 | 1.45E-06 |
| COPA | Coatomer subunit alpha;Xenin;Proxenin | 3.65E-06 | 1.13E-06 |
| DSG2 | Desmoglein-2 | 3.65E-06 | 8.73E-07 |
| DBN1 | Drebrin | 3.65E-06 | 1.03E-06 |
| RPL12 | 60S ribosomal protein L12 | 3.97E-06 | 1.94E-06 |
| H3F3B | Histone H3;Histone H3.2;Histone H3.1t;Histone H3.3;Histone H3.1;Histone H3.3C | 3.97E-06 | 1.45E-06 |
| LETM1 | LETM1 and EF-hand domain-containing protein 1, mitochondrial | 4.02E-06 | 1.51E-06 |
| MYH10 | Myosin-10 | 4.86E-06 | 1.34E-06 |
| GNB2 | Guanine nucleotide-binding protein G(I)/G(S)/G(T) subunit beta-2;Guanine nucleotide-binding protein subunit beta-4 | 5.1E-06 | 2.12E-06 |
| DNAJB11 | DnaJ homolog subfamily B member 11 | 5.41E-06 | 1.94E-06 |
| ATP5O | ATP synthase subunit O, mitochondrial | 7.39E-06 | 2.64E-06 |
| PHGDH | D-3-phosphoglycerate dehydrogenase | 8.22E-06 | 2.72E-06 |
| TBL2 | Transducin beta-like protein 2 | 2.35E-05 | 8.02E-06 |
| WDR6 | WD repeat-containing protein 6 | 2.89E-05 | 1.08E-05 |

**Supplementary Table S1. List of potential proteins which interact with DP fusion protein based on proteomic analysis.**

Supplementary Table S2.

| **Pathway Term** | **Description** | **p-value** | **FDR** |
| --- | --- | --- | --- |
| Metabolism of proteins Homo sapiens R-HSA-392499 | Metabolism of proteins Homo sapiens R-HSA-392499 | 3.012E-06 | 0.005 |
| double-stranded RNA binding (GO:0003725) | double-stranded RNA binding (GO:0003725) | 6.8601E-05 | 0.007 |
| tRNA binding (GO:0000049) | tRNA binding (GO:0000049) | 6.6877E-05 | 0.007 |
| mRNA binding (GO:0003729) | mRNA binding (GO:0003729) | 7.1708E-05 | 0.007 |
| translation factor activity, RNA binding (GO:0008135) | translation factor activity, RNA binding (GO:0008135) | 6.3133E-05 | 0.007 |
| single-stranded RNA binding (GO:0003727) | single-stranded RNA binding (GO:0003727) | 6.5609E-05 | 0.007 |
| snoRNA binding (GO:0030515) | snoRNA binding (GO:0030515) | 6.1527E-05 | 0.008 |
| rRNA binding (GO:0019843) | rRNA binding (GO:0019843) | 6.1131E-05 | 0.008 |
| telomerase RNA binding (GO:0070034) | telomerase RNA binding (GO:0070034) | 6.1131E-05 | 0.008 |
| primary miRNA binding (GO:0070878) | primary miRNA binding (GO:0070878) | 6.0737E-05 | 0.008 |
| siRNA binding (GO:0035197) | siRNA binding (GO:0035197) | 6.0345E-05 | 0.009 |
| AU-rich element binding (GO:0017091) | AU-rich element binding (GO:0017091) | 6.0345E-05 | 0.009 |
| miRNA binding (GO:0035198) | miRNA binding (GO:0035198) | 6.0345E-05 | 0.009 |
| piRNA binding (GO:0034584) | piRNA binding (GO:0034584) | 5.9956E-05 | 0.01 |
| snRNA binding (GO:0017069) | snRNA binding (GO:0017069) | 5.9956E-05 | 0.01 |
| G-quadruplex RNA binding (GO:0002151) | G-quadruplex RNA binding (GO:0002151) | 5.9568E-05 | 0.01 |
| pre-miRNA binding (GO:0070883) | pre-miRNA binding (GO:0070883) | 5.9568E-05 | 0.01 |
| RNA cap binding (GO:0000339) | RNA cap binding (GO:0000339) | 5.9568E-05 | 0.015 |
| RNA stem-loop binding (GO:0035613) | RNA stem-loop binding (GO:0035613) | 5.9183E-05 | 0.01 |
| pre-mRNA binding (GO:0036002) | pre-mRNA binding (GO:0036002) | 5.9183E-05 | 0.01 |
| histone pre-mRNA DCP binding (GO:0071208) | histone pre-mRNA DCP binding (GO:0071208) | 5.9183E-05 | 0.01 |
| BRE binding (GO:0042835) | BRE binding (GO:0042835) | 5.9183E-05 | 0.02 |
| telomeric repeat-containing RNA binding (GO:0061752) | telomeric repeat-containing RNA binding (GO:0061752) | 5.9183E-05 | 0.02 |
| 7S RNA binding (GO:0008312) | 7S RNA binding (GO:0008312) | 5.9183E-05 | 0.02 |
| 21U-RNA binding (GO:0034583) | 21U-RNA binding (GO:0034583) | 5.88E-05 | 0.02 |
| misfolded RNA binding (GO:0034336) | misfolded RNA binding (GO:0034336) | 5.88E-05 | 0.02 |
| base pairing with RNA (GO:0000498) | base pairing with RNA (GO:0000498) | 5.88E-05 | 0.03 |
| actin filament binding (GO:0051015) | actin filament binding (GO:0051015) | 0.00028692 | 0.03 |
| ribonuclease P RNA binding (GO:0033204) | ribonuclease P RNA binding (GO:0033204) | 5.88E-05 | 0.03 |
| N6-methyladenosine-containing RNA binding (GO:1990247) | N6-methyladenosine-containing RNA binding (GO:1990247) | 5.88E-05 | 0.04 |
| alpha-aminoacyl-tRNA binding (GO:1904678) | alpha-aminoacyl-tRNA binding (GO:1904678) | 5.88E-05 | 0.05 |
| actin lateral binding (GO:0003786) | actin lateral binding (GO:0003786) | 0.00063334 | 0.06 |
| RNA binding (GO:0003723) | RNA binding (GO:0003723) | 5.88E-05 | 0.06 |

**Supplementary Table S2.** **Gene enrichment pathway analysis based on proteomic data.** Ranked pathways of significance based on differential protein precipitation in response to PKA-Cα immunoprecipitation in A9 and A11 compared to HEK-WT.
